# Identifying Neural Biomarkers of Risk-Taking from Intracranial EEG Recordings

**DOI:** 10.1101/2025.10.08.681279

**Authors:** Yuxin Guo, Amanda Merkley, Stephen Jaffee, Alexander C. Whiting, Pulkit Grover

**Author notes:** These authors contributed equally (Yuxin Guo, Amanda Merkley).

## Abstract

Risk-taking behavior is associated with neuropsychiatric disorders such as addiction. In this study, we present a data-driven approach to identify neural biomarkers of risk-taking from intracranial stereo-electroencephalography (sEEG) recordings. Using time and frequency domain features, we train a classifier to distinguish between risk-taking and risk-avoidance behaviors. Based on data from a single patient, the model achieves an accuracy of 68.5%, which is significantly above chance. These results highlight the potential of identifying risk-taking states from invasive recordings.

**Clinical Significance:** This work demonstrates the feasibility of identifying risk-related biomarkers from intracranial recordings, highlighting a promising direction for closed-loop neuromodulation for neuropsychiatric conditions.

## I. INTRODUCTION

Understanding risk-taking behaviors is crucial, as abnormal risk-taking is associated with several neuropsychiatric conditions, such as addiction and bipolar disorder [1]– [3]. Consequently, there is growing interest in identifying the neural biomarkers underlying risk-taking behaviors to support diagnosis, monitoring, and intervention, potentially enabling closed-loop neural stimulation strategies upon detection of such biomarkers. In this work, we record intracranial stereoelectroencephalography (sEEG) signals from a participant engaged in a task that involves voluntary risk-taking. We then use the resulting data to obtain biomarkers of risk-taking.

Specifically, the participant engaged in the Balloon Analogue Risk Task (BART). BART has been demonstrated to assess participants’ propensity for risk-taking in laboratory settings [4], and has been used to study real-world risk-related behaviors such as substance-use [5], [6] and gambling [7]. During BART, participants are sequentially presented with several virtual balloons on a laptop screen. For each balloon, participants decide at each step whether to inflate the balloon—which either increases the potential reward or causes the balloon to explode—or to cash in and secure the cumulative reward (as shown in Fig. 1). If a balloon explodes (when the participant chooses to inflate), which occurs at a random time unknown to the participant, all accumulated reward from that balloon is lost. Participants thus need to make decisions under uncertainty and balance between inflation for higher reward (risk-taking) and cashingin to avoid loss (risk-avoidance).

**Fig. 1.**
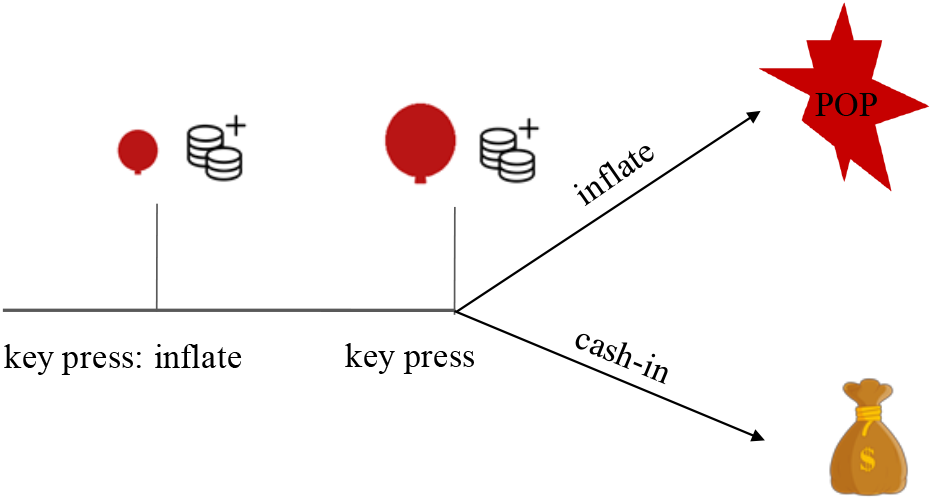
Illustration of BART. At each step, the participant chooses whether to inflate the balloon—thereby increasing potential reward at the risk of explosion—or to cash in and secure the accumulated reward. An explosion results in the loss of all reward from that balloon.

Previous studies have used functional magnetic resonance imaging (fMRI) to investigate the neural correlates of voluntary risk-taking during BART. Rao et al. [8] observe that the mesolimbic-frontal regions (e.g. anterior insula, anterior cingulate, and medial frontal cortex) are active during volitional risk-taking. Fukunaga et al. [9] observed increased anterior cingulate and inferior frontal gyrus activation when participants discontinued balloon inflation, consistent with loss aversion signaling. In this work, we analyzed sEEG recordings, i.e., invasive brain recordings, providing us direct access to several deep brain regions, with a high spatiotemporal resolution that is not achieavable by noninvasive methods.

We formulate the biomarker identification problem as a binary classification task, where the goal is to predict behavior (risk-taking vs. risk-avoidance) from sEEG recordings. Here, we define candidate biomarkers as time- and frequency-domain features derived from sEEG signals. The choice of these features is informed by the NeuroPace Responsive Neurostimulation (RNS) system [10], which could in the future enable programming in closed-loop stimulation systems. Using these features, we develop a classification model that achieves above-chance accuracy (i.e. 68.5%) in distinguishing risk-taking from risk-avoidance behaviors.

### II. Problem Statement

We formulate behavior decoding as a binary classification task: risk-taking versus risk-avoidance. Considering the clinical application of closed-loop stimulation, we constrain our analysis to features derived from at most two recording channels, in line with typical limitations of implantable neuromodulation devices.

We make the following assumptions to define behavioral labels: for balloons where the participant ultimately chose to cash in, we assume that the final key-press (cash-in) reflects risk-avoidance, while the immediately preceding inflation key-press reflects volitional risk-taking (see Fig. 2). Only the immediately preceding inflation key is considered because we assume that it carries the highest risk in each trial. These two behavioral events serve as the labels for supervised classification, and model-inputs are derived from the neural activity preceding each event.

**Fig. 2.**
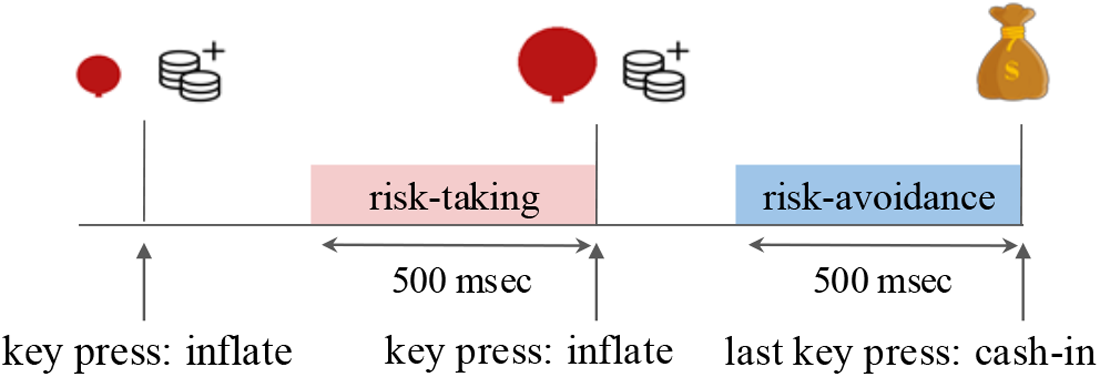
For each balloon in which the participant ultimately cashed in, the final key press (cash-in) is labeled as risk-avoidance, while the second-to-last key press (inflate) is labeled as risk-taking.

### III. Methods

#### A. Data Acquisition

The experiment was conducted with a 36-year-old male patient undergoing clinical sEEG monitoring for seizure localization at Allegheny Health Network (AHN). All procedures were approved by the AHN Institutional Review Board. sEEG probes were placed bilaterally, covering deep brain regions including the amygdala, hippocampus, insula, gyrus rectus, etc. The patient performed BART on a laptop.Behavioral key presses were recorded and synchronized with sEEG recordings using event triggers via the Natus Quantum® LTM Amplifier recording system (Natus Medical Inc), sampled at 1024 Hz. The risk-taking (balloon inflation) events were marked by pressing the space bar. On the other hand, risk-avoidance (cash-in) events were marked by pressing the enter key. A total of 90 balloons (including both cash-in and explosion trials) were completed by the participant within 15 minutes. The sEEG recordings analyzed in this study were collected during clinically stable periods outside of seizure events.

#### B. Data Pre-processing

For noise removal, signals were band-pass filtered (0.5-100 Hz) and notch filtered at 60 Hz to remove line noise. To remove shared global artifacts, we performed bipolar rereferencing by subtracting signals from adjacent contacts along each probe [11]. Among all contacts on the recording probes, we selected those located within brain regions previously associated with risk-taking behaviors. These regions include the orbitofrontal cortex [12], anterior cingulate cortex [13], amygdala [14], parahippocampal gyrus [15], and insula [16], resulting in 34 bipolar channels. Based on the labeling strategy described in Section II, we identified all balloons for which the participant chose to cash in, and extracted 500 ms data segments preceding both the risk-taking and risk-avoidance events (as illustrated in Fig. 2). Due to the length of the inter-click interval, only 500 ms is taken for analysis to prevent overlapping with neighboring key presses. This yielded a balanced dataset of 58 data points, where each data point has recordings from 34 bipolar channels.

#### C. Biomarker Determination

As described in Section II, our goal is to identify time and frequency domain biomarkers from only two recording channels, in accordance with the constraints of closed-loop RNS devices. Our approach involves two steps (Fig. 3). First, we perform channel selection using a majority-vote procedure inspired by the idea of stability selection [17], where the most frequently top-ranked channels are chosen across randomized subsets of data (see Section. III-C.2). This step is conducted using Dataset 1 (40% of the data, randomly selected). Then, a classifier is trained and evaluated on Dataset 2 (the remaining data) using features extracted from the selected channels.

**Fig. 3.**
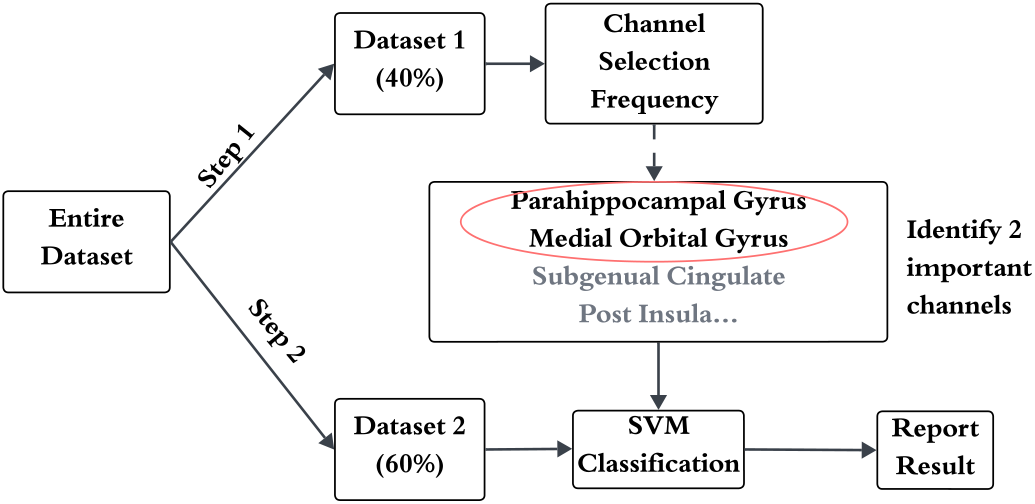
Overview of the two-step analysis pipeline. The dataset is split into two subsets: Dataset 1 is used for channel selection using a stability-inspired majority-vote procedure; Dataset 2 is then used to train and evaluate an SVM classifier on features extracted from the two most frequently selected channels.

##### 1) Feature Extraction

From each channel’s raw signal *X*_*c*_ for *c* = 1, …, 34, we extract time- and frequency-domain features. Specifically, we compute:

- **Band power features:** Power is computed in six frequency bands—delta (1-4 Hz), theta (4-8 Hz), alpha (8-12 Hz), low beta (12-20 Hz), high beta (20-30 Hz), and gamma (30-70 Hz).
- **Line length (LL):** Measures the average change in signals across time. Let *τ* be the window length in seconds, *f*_*s*_ the sampling rate in Hz, and *K* = *τ f*_*s*_ the total number of samples in that window. Then for channel *c* we define:

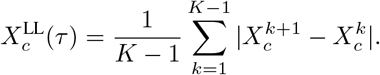
- **Area under the curve (AUC):** Related to signal energy, defined as:

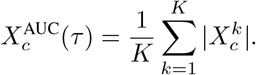

The LL and AUC features are informed by NeuroPace RNS system’s built-in detectors. To capture both long- and short-term dynamics, we compute 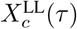 and 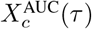 for *τ* = 0.5 s (i.e. over the entire time window) and *τ* = 0.25 s (i.e. over the last 250ms). This results in two 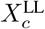 values and two 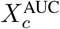 values per channel. Combined with the six band power features, each channel is represented by a total of 10 features.

##### 2) Channel Selection

With the goal of identifying the most informative channels, we looked for channels that consistently and reliably predict behavior labels despite variations in the training split (i.e., random training subsets). As shown in Algorithm 1, we randomly split Dataset 1 into training and test sets. For each split *n*, a support vector machine (SVM) with a radial basis function (RBF) kernel was trained using only the features from channel *c*. The model’s regularization parameter *λ* was tuned via grid search over the values *λ* ∈ {0.1, 1, 10, 100} using 3-fold cross-validation on the training set. The resulting model was then evaluated on the hold-out test set to compute an accuracy score *a*_*c*_.

This process is repeated for *N* = 500 different random splits of Dataset 1. For each split, we identify the channel *c* that achieves the highest accuracy *a*_*c*_, for the specific training subset, and record it as a “winner.” After *N* repetitions, we record how often each channel was selected as the top performer. The two channels with the highest selection counts (*c*^(1)^ and *c*^(2)^) are retained as the most informative channels for the next step of analysis.

##### 3) Model Training and Evaluation

Jointly using the two selected channels *ĉ*^(1)^ and *ĉ*^(2)^, we evaluated classification performance on Dataset 2. We repeated the following procedure *M* = 100 times: randomly split Dataset 2 into 70% training and 30% testing subsets (stratified), extract features corresponding to channels *ĉ*^(1)^ and *ĉ*^(2)^, train an SVM classifier with 3-fold cross-validation on the training set, and compute the classification accuracy on the test set. We report the mean and standard deviation of the accuracy across all *M* repetitions to obtain average performance over random splits of the data.

#### D. Spectrogram Analysis

To analyze the temporal dynamics of spectral power associated with each behavioral event (i.e., risk-taking and risk-avoidance), we computed time–frequency spectrograms from sEEG signals. On each 500 ms data segment (extracted as described in Fig. 2) and each recording channel, we applied the short-time Fourier transform (STFT) using a 128-sample Hamming window (125 ms), 50% overlap, and a sampling rate of 1024 Hz.

Let 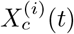 denote the signal from channel *c* at data segment *i*. For each segment and channel, the spectrogram 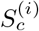 was computed. We then computed the average spectrogram across all data segments in each condition (risk-taking and risk-avoidance) for each channel:

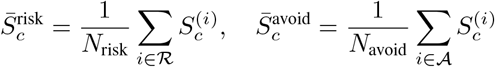

where ℛ and *A* denote the sets of risk-taking and risk-avoidance signal segments, respectively.

##### Algorithm 1

Channel Selection via Majority Vote

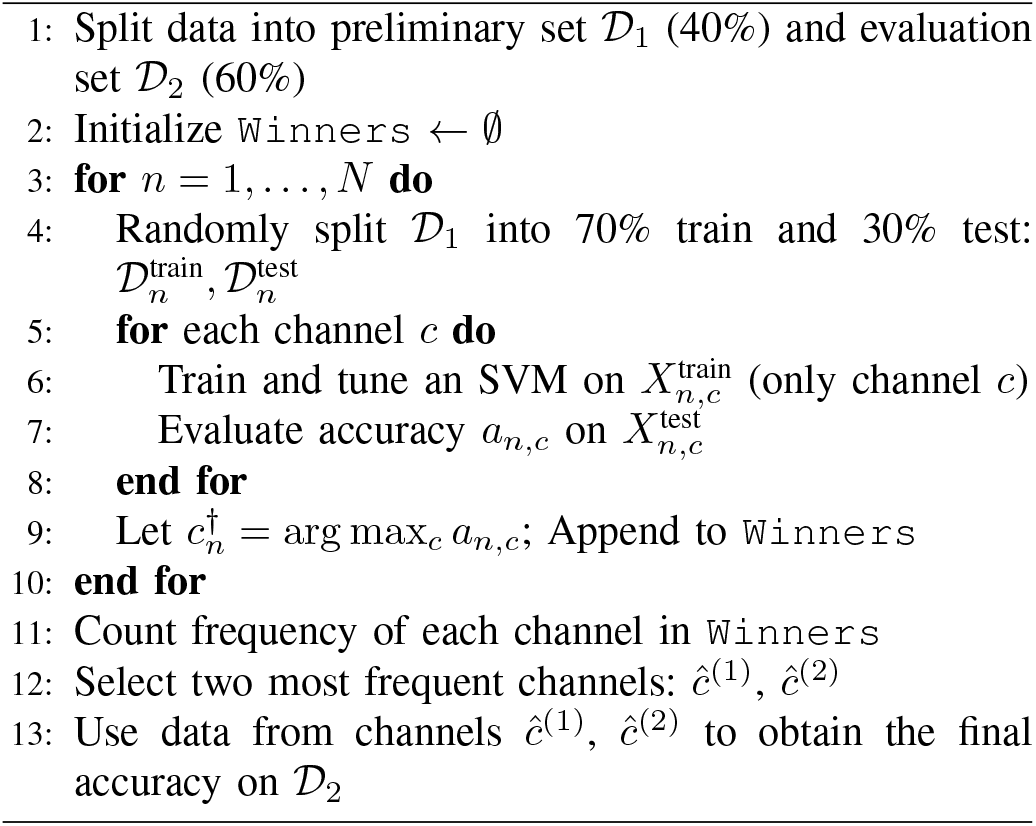

To compare spectral dynamics between conditions, we computed the relative difference spectrogram as:

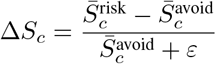

with a small constant *ε* = 10^*-*12^ added for numerical stability. ∆*S*_*c*_ quantifies the ratio of difference between risk-taking and risk-avoidance conditions for each channel.

## IV. Results

### A. Channel Selection Results

Using the procedure described in Section III-C.2, the two most frequently selected channels were located in the parahippocampal gyrus (PHG) and medial orbital gyrus (MOG). Among the 34 available channels, the PHG channel and MOG channel were selected with the highest counts—73 and 70 times, respectively—across *N* = 500 random training subsets. Figure 4 shows the selection counts of the top 10 channels. Notably, there is a sharp drop in selection counts after the third most frequently selected channel (another MOG channel, lMOG2), suggesting diminishing returns beyond the top few. Additionally, the third-ranked channel is located one contact away from the second, and likely shares correlated information due to spatial proximity. This supports that selecting the top two channels captures the most informative signals. This finding is consistent with prior work implicating both MOG and PHG in risk-related decisionmaking and cognitive control under uncertainty [8], [12], [18].

**Fig. 4.**
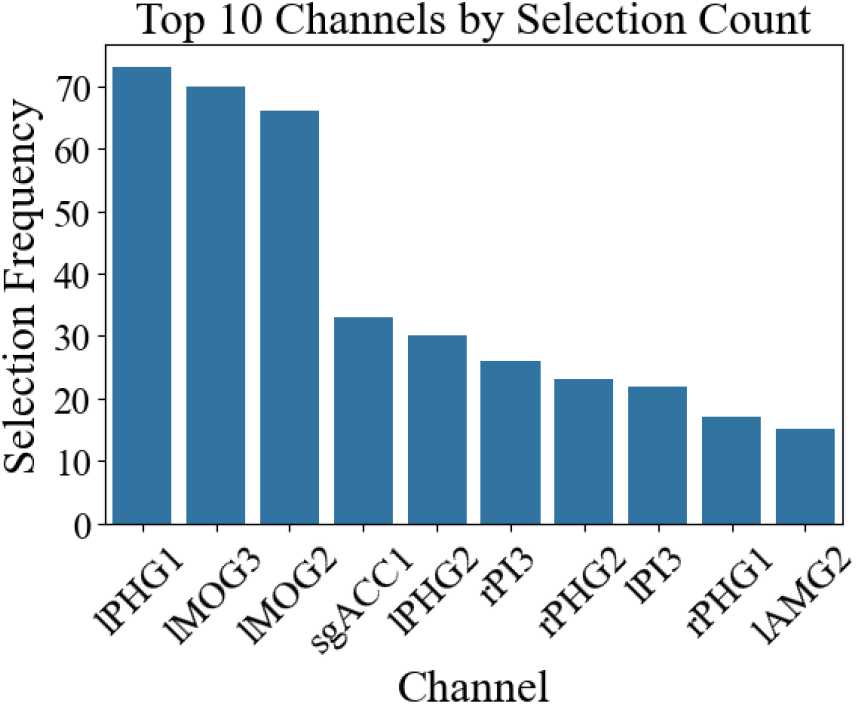
Top 10 channels ranked by selection counts across *N* = 500 random training subsets. Channel labels follow the format: hemisphere + region + contact number. ‘l’ and ‘r’ indicate the left and right hemisphere, respectively; the numeric suffix denotes contact depth (e.g., lPHG1 = left parahippocampal gyrus, contact 1 from the distal end of the probe). Region abbreviations: PHG = parahippocampal gyrus, MOG = medial orbital gyrus, sgACC = subgenual cingulate, PI = posterior insula, AMG = amygdala.

To further characterize the neural dynamics of these regions, we analyzed time–frequency spectrograms comparing risk-taking and risk-avoidance events (see Section III-D). As shown in Fig. 5, the MOG channel shows elevated beta-band activity during risk-taking, with a maximum relative increase of 77% (1.77×) at 27 Hz, approximately 0.31 seconds before the final key press. The PHG channel (Fig. 6) showed a 72% increase in theta-band power (1.72×) during risk-taking, peaking at 5 Hz and 0.31 seconds before the final key press.

**Fig. 5.**
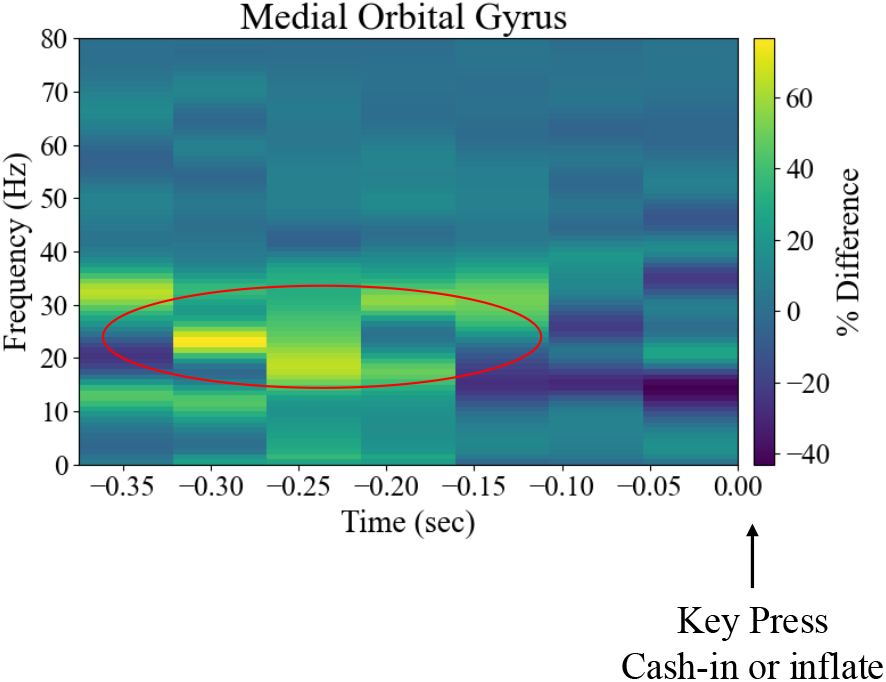
Relative power difference spectrogram for medial orbital gyrus.

**Fig. 6.**
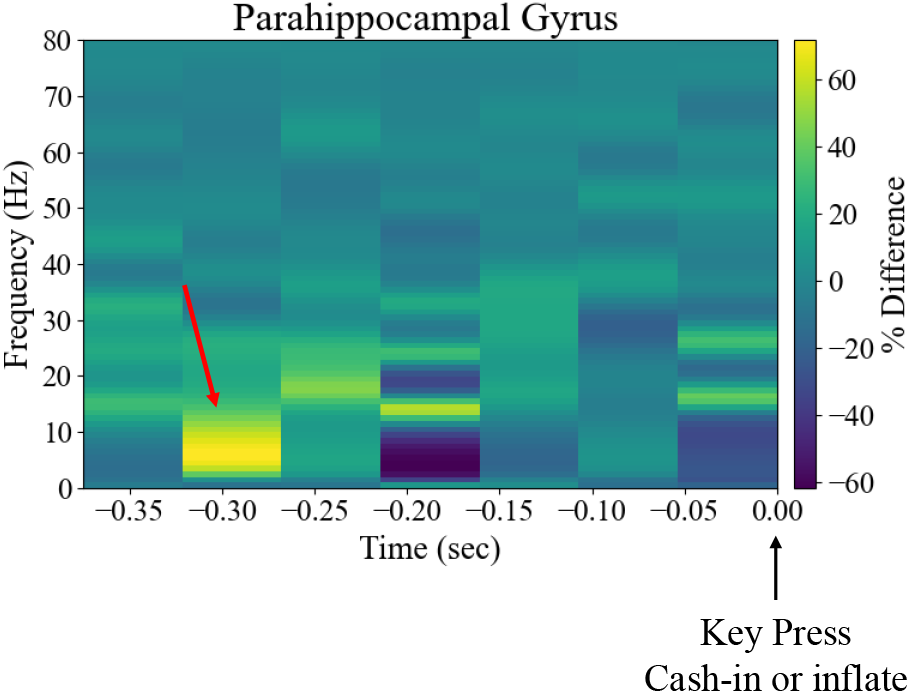
Relative power difference spectrogram for parahippocampal gyrus.

### B. Classification Results

We trained an SVM using features from the two most informative channels (i.e. PHG and MOG) on Dataset 2. To account for variability introduced by random train/test splits, the training and evaluation process was repeated 100 times (see Section. III-C.3). The model achieved a mean classification accuracy of 68.5% *±* 9.1%. To assess whether this performance was significantly above chance (where chance is 50% accuracy), we conducted a permutation test with 20,000 label shuffles. The null hypothesis is that the labels are statistically independent of the features, and the test statistic was defined as the mean accuracy across repetitions. The resulting *p*-value was computed as 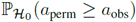, where *a*_perm_ and *a*_obs_ denote the mean classification accuracies under the null (permuted labels) and observed data, respectively. This yielded a *p*-value of 0.0039, indicating statistically significant above-chance performance.

Table I summarizes the performance of the proposed 2-channel model (overall accuracy, true positive rate, and false positive rate) and compares it to an SVM trained on features from all 34 channels (without channel selection). Despite using features from much fewer channels, the 2-channel model achieves higher accuracy and a lower false positive rate. These results suggest that reliable biomarker identification requires rejecting noisy channels, as this improved performance may be due to reduced overfitting and the elimination of noisy or less informative features/channels.

**TABLE I.**
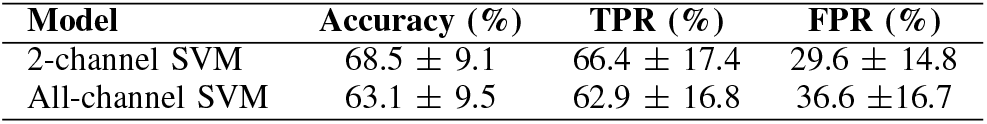
Performance of SVM Classifiers.

## V. Conclusions and Discussions

We present a data-driven approach for identifying risk-related neural biomarkers from intracranial sEEG recordings collected during BART. Using behavioral labels (risk-taking vs. risk-avoidance), we demonstrated that above-chance accuracies can be achieved using features from just two channels, specifically, the parahippocampal gyrus and medial orbital gyrus. Both regions have been previously implicated in decision-making and cognitive control under uncertainty [8].

### Lack of generalizability to other participants

This study was carried out with one patient, which limits the generalizability of the identified biomarkers. The selected channels and features for decoding risk-taking behavior may vary across individuals due to anatomical and functional differences. Future work should validate these findings across a larger cohort.

### Aligning Models with RNS Device’s Hardware Limitations

Additionally, while the features used in this work are compatible with those supported by the NeuroPace RNS system, the classification model (i.e. SVM with an RBF kernel) does not reflect the logic used by the device’s built-in detectors, which relies on threshold-based rules and logical operators. To enable direct programming of the RNS device, future efforts should optimize within the NeuroPace-compatible hypothesis class and develop detection rules aligned with device constraints.

### Task Generalizability

While BART is a well-validated task for assessing risk-taking propensity in laboratory settings, it remains an approximation for real-world behaviors such as substance use addiction. The neural biomarkers identified in this study are specific to BART-driven risk-taking behaviors and may not directly translate to naturally occurring behaviors. Bridging this gap will require future studies that incorporate real-world risk-taking data.

### Toward Causal Interventions

While this study demonstrates a correlation between biomarkers and risk-taking behavior, it does not establish a causal relationship. Future work is needed to test whether targeted stimulation can suppress these biomarkers and subsequently modulate behavior. Even if a causal link is validated, identifying optimal stimulation parameters, including the anatomical target and waveform characteristics, will be essential for designing effective closed-loop neuromodulation interventions.

## ACKNOWLEDGMENT

This work was supported by a fellowship from the Center for Machine-Learning and Health (Y.G.), and a fellowship from the NSF Graduate Research Fellowship Program (A.M).

## Notes

### Competing Interest Statement

The authors have declared no competing interest.

